# Toward D-peptide biosynthesis: Elongation Factor P enables ribosomal incorporation of consecutive D-amino acids

**DOI:** 10.1101/125930

**Authors:** Po-Yi Huang, Fanny Wang, Kamesh Narasimhan, Kelly Chatman, John Aach, Sunia A. Trauger, Ryan Spoering, George M. Church

**Affiliations:** Department of Genetics, Harvard Medical School, Boston, Massachusetts, USA; Virginia Tech-Wake Forest University School of Biomedical Engineering and Sciences, Wake Forest University School of Medicine, Winston-Salem, North Carolina, USA; FAS Small Molecule Mass Spectrometry Facility, Harvard University, Boston, MA, USA; Lab Machinist Solutions, Waltham, Massachusetts

## Abstract

To maintain stereospecific biochemistry in cells, living organisms have evolved mechanisms to exclude D-amino acids (*^D^*AA) in their protein synthesis machinery, which also limits our exploration of the realm of mirror-image molecules. Here, we show that high affinity between EF-Tu and aminoacyl-tRNA promotes D-amino acid incorporation. More strikingly, Elongation Factor P efficiently resolves peptidyl transferase stalling between two consecutive D-amino acids, and hence enables the translation of D-peptides.

---

Life is an anti-entropic phenomenon with two mutually-reinforcing characters: homochirality and stereospecific catalysis. The exclusive presence of L-amino acids in proteins of the living world is a prominent example of this. However, D-amino acid containing peptides (DAACP) are still present in microbial, fungal and amphibian secretions, and often carry interesting bioactivities^1^. In nature, these molecules are made through non-ribosomal pathways, such as non-ribosomal peptide synthesis or post-translational modification, e.g. epimerization. DAACPs have been shown to have prolonged half lives in serum and resistance toward proteases without immunogenicity^2^, which is a desirable property in therapeutic reagents. Therefore, by gradually rewiring the present bio-machineries to overcome natural *^D^*AA barrier, we aim to build a bridge leading us to the space of mirror-image biomolecules, where they can serve as new tools for the biotech and pharmaceutical industry.

Core protein translation machinery exhibits significant discrimination of *^D^*AAs from L-amino acids (*^L^*AAs) at three steps^3^: aminoacylation of tRNAs by aminoacyl-tRNA synthetases (aaRSes), formation of ternary complexes with EF-Tu-GTP, and peptide-bond formation catalyzed by the ribosome (Figure 1a). In order to study these discriminations without the interference from D-amino acid oxidase and D-aminoacyl-tRNA deacylase^4,5^, we use a purified *E. coli* protein synthesis system (PURE)^6^, along with chemically acylated D-aminoacyl-tRNA^7^, as a model system for engineering ^*D*^AA-tolerant translation machinery. We adapted the amber codon read through assay described in Fujino *et al.*^8^ with slight modifications to assess *^D^*AA incorporation during elongation (Figure 1b). In brief, mRNA templates encoding an N-terminus FLAG epitope and a C-terminus stretch of artificial peptide composed of six different amino acids, either with or without an amber stop codon (UAG) in between, are translated *in vitro* with purified ribosomes, translational factors and aaRSes, in the presence or absence of amber codon suppressor tRNAs that have been chemically acylated with *^D^*AAs or *^L^*AAs. The peptides produced were resolved in denaturing PAGE gels and visualized by western blotting with anti-FLAG antibodies. On examining the incorporation of various *^L^*AA, *^D^*AA, and an α, α-dialkylamino acid with the orthogonal amber suppressor AsnE2 tRNA_CUA_^8^, only L-valine yielded measurable incorporation, while L-alanine and all others tested amino acids could not be detected (Figure 1c), suggesting the limited sensitivity of western blotting. This could be explained by the higher affinity of EF-Tu to *^L^*Val-tRNA^AsnE2^ than to *^L^*Ala-tRNA^AsnE2^ with the estimated dissociation constants (K_D_) of 88.3 nM and 532 nM respectively, while the K_D_ of 20 *^L^*AA-tRNA bodies are between 5.7 to 92.0 nM (Supplementary Table 1)^9^. Since the amount of all tRNA used in typical PURE translation (13-53 μM) is greater than that of EF-Tu (2-10 μM, Supplementary Table 2), the sequestering of EF-Tu could in theory limit EF-Tu access to chemically acylated *^L/D^*AA-tRNAs, especially for *^D^*AA, since *^D^*Tyr acylated tRNA has been shown to result in 25 fold decrease than *^L^*Tyr in affinity toward EF-Tu^10^. In addition, the fast kinetics of aminoacyl hydrolysis when not shielded by EF-Tu^10^ and the disadvantage that they cannot be regenerated by aaRSes like their competing normal *^L^*AA-tRNA pairs, all lead to poor *^D^*AA incorporation yield.

**Figure 1.**
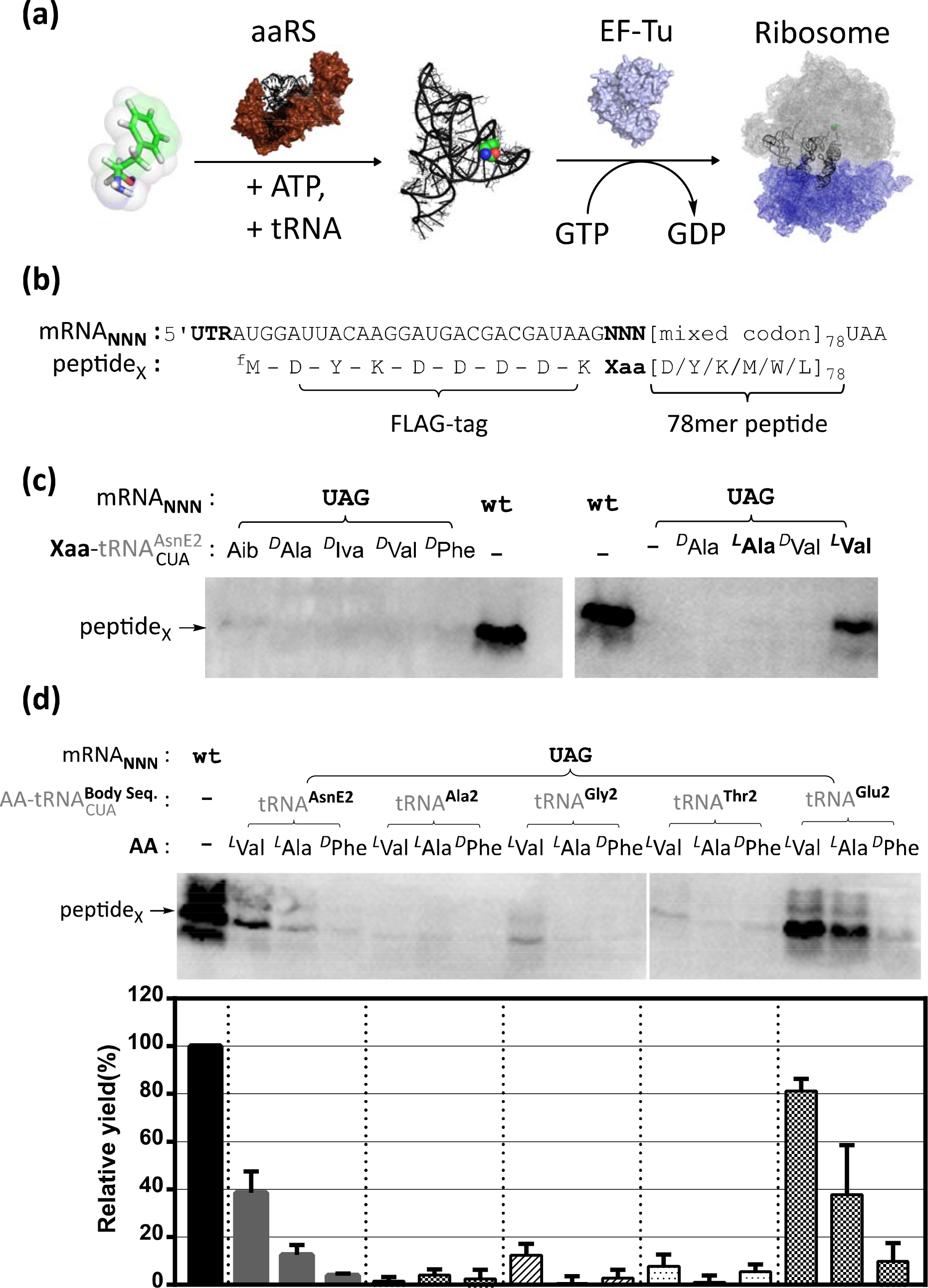
(a) Three steps blocking *^D^*AA in protein synthesis: aaRS bias to *^L^*AA acylation, EF-Tu bias on binding to *^L^*AA-tRNA and ribosome bias in catalyzing *^L^*AA peptidyl transfer. (b) Protein translation read-through assay set up. mRNA templates encoding N-terminus FLAG epitope and a C-terminus stretch of artificial peptide composed of six different amino acids, either with or without an amber codon in between. *In vitro* translated peptides are resolved in denaturing PAGE gel and visualized by western blotting with anti-FLAG antibody. (c) Western Blot of *^L/D^*AA read-through assay carried by AsnE2-tRNA body. Only *^L^*Val yields measurable incorporation. 12 μM of chemical acylated AA-tRNA and 2 μM of EF-Tu are used. mRNA with label “wt” is a template without middle NNN codon and hence its synthesis doesn’t require tRNA_CUA_. (d) Various tRNA backbones with CUA anticodon are tested for the incorporation of *^L^*val, *^L^*Ala and *^D^*Phe, under 24 μM of chemical acylated AA-tRNA and 25 μM of EF-Tu. tRNA^Glu2^ shows superior incorporation of the AAs tested. The signal intensities are normalized against the signal of “wt” peptide, and the error bar represents the standard deviation of each triplicated experiment.

Incorporation could possibly be improved by additional EF-Tu and higher concentrations of charged tRNAs in the *in vitro* translation system, but we had earlier found that this only achieved a moderate improvement in *^L^*Ala^11^. We therefore thought to compensate for the disadvantage of low-incorporating *^L/D^*AAs by using a tRNA body that binds very efficiently to EF-Tu, noting that the K_D_s of tRNA bodies to EF-Tu-GTP span 600-fold from the weakest to the strongest^9^. Therefore, we chose four tRNA bodies (Glu2, Thr2, Gly2 and Ala2, sequences provided in Supplementary Methods) which exhibited strong affinities toward EF-Tu-GTP to test whether they could promote incorporation of weak EF-Tu binding *^L/D^*AAs into peptides. This approach has recently been reported by Achenbach *et al*.^12^, in which they found better incorporation with tRNA^Gly^ than tRNA^Tyr^ when both are acylated with same *^D^*AAs^12^.We found that the Glu2 tRNA body outperformed Thr2, Ala2, Gly2 and AsnE2 tRNA body in *^L^*Ala incorporation, and also yielded ca. 14% *^D^*Phe incorporation, relative to peptides synthesized without amber codon read through (noted as ‘wt’ in all the figures here) (Figure 1d). Here, the absence of RNA base modifications in the *in vitro* transcribed tRNAs could have affected their kinetics in the ribosome^13^.

Although many groups^8,12,14,15^ have now shown that single *^D^*AAs can be incorporated by the ribosome, *^D^*AA-*^D^*AA consecutive incorporations are generally barely detectable and show greatly reduced kinetics, suggesting that ribosomal catalytic mechanisms are highly compromised. We had previously attempted to mutagenize the peptidyl transferase center (PTC) and select for improved *^D^*AA incorporation, but this proved unsuccessful as none of the surviving mutants exhibited enhanced protein synthesis capacity regardless of whether *^D^*AA were encoded in the template or not^11^. Recently, elongation factor P (EF-P) has been reported to resolve ribosome stalling upon incorporating consecutive polyprolines^16,17^. eIF-5A, the eukaryotic homolog of bacterial EF-P, has also been shown to help the ribosome resolve polyproline-stalling by orienting the CCA-3’ end of the P-site tRNA with a nearby rRNA uL^16^ loop in a productive conformation^18^. Since EF-P is also found to facilitate translation initiation^19^, i.e. peptidyl transfer of P-site formylmethionyl-tRNA, this mode of action might operate more generally. Hence we examined whether EF-P might improve *^D^*AA read-through in our *in vitro* system. To our surprise, EF-P (2 μM, the concentration used to resolve polyproline stalling ^19^) not only promoted single *^D^*Phe incorporation, but also improved consecutive *^D^*Phe-*^D^*Phe read-through event up to 10% relative yield at 4 µM (Figure 2a). EF-P’s effect for *^D^*AA diminished at concentrations above 8 μM, but interestingly, EF-P does not show any inhibitory effect on ‘wt’ peptide translation at this high concentration range (Figure 2b).

**Figure 2.**
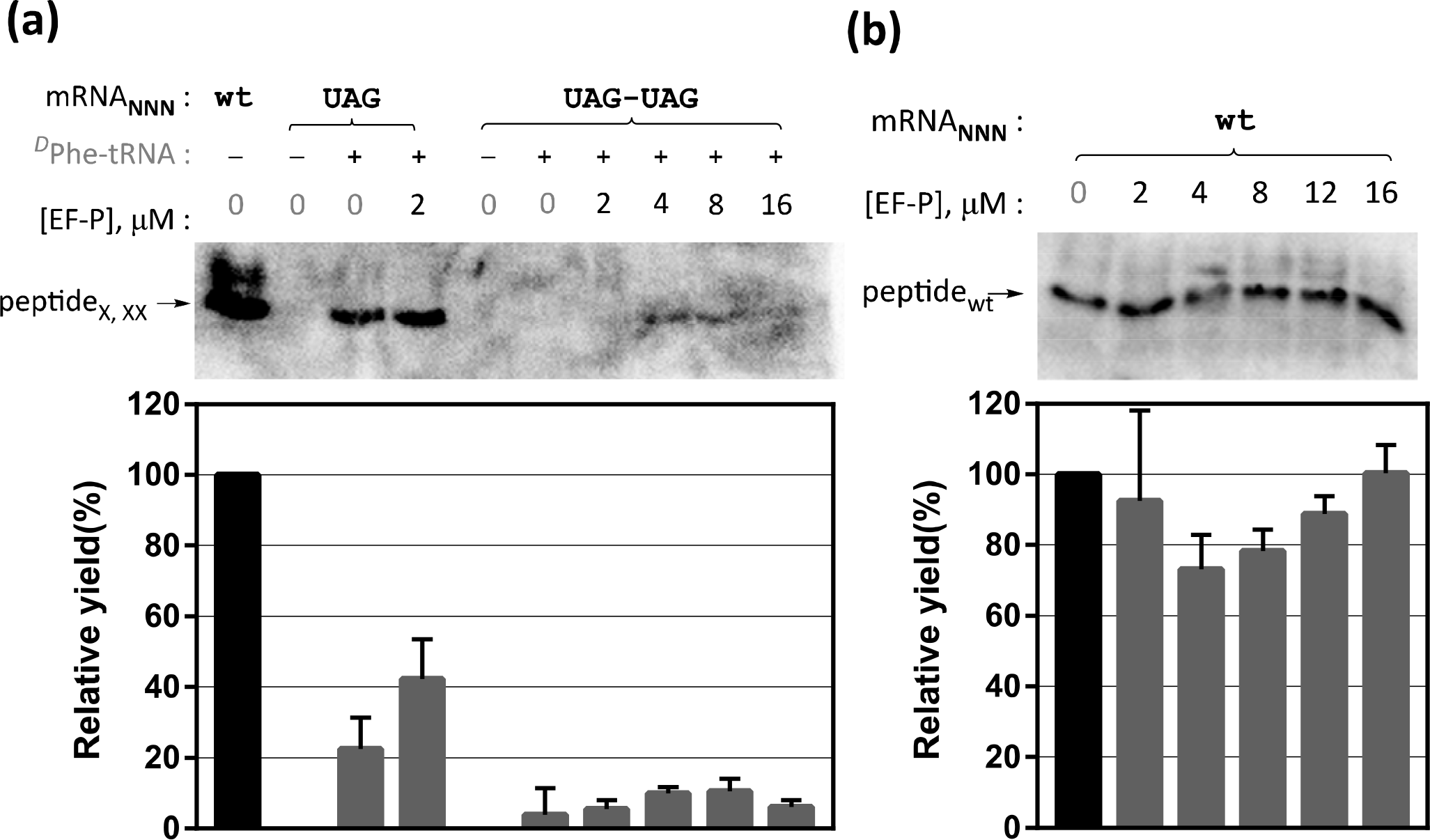
(a) Effect of EF-P on *^D^*Phe incorporation. At 2 μM of EF-P, yield of single incorporation is promoted by 1.9 times; at 4 μM and 8 μM, EF-P promotes consecutive *^D^*Phe incorporation up to 10% relative yield, however, this facilitating effect reduced at 16 μM of EF-P. (b) Effect of EF-P on ‘wt’ peptide synthesis. EF-P shows slight inhibitory effect at 4 μM, but has less effect at higher concentration.

Given that substantial consecutive *^D^*AA incorporation could be achieved with EF-P, the next question was how long a string of *^D^*AAs the ribosome could potentially polymerize. Since chemically acylated *^D^*AA-tRNAs suffer from fast hydrolysis at physiological pH, we sought to move to a system in which *^D^*AA-acylated tRNAs could be maintained by an aaRS. Tyrosyl-tRNA synthetase has been long known to support *^D^*Tyr charging, both in bacteria and in yeast^10,20^. We picked wild type *M. jannaschii* tyrosyl-tRNA synthetase (*Mj*-YRS) because its determinant is simple, i.e. G2:C72^21^, and could be transplanted into tRNA^Glu2^ without compromising its strong affinity to EF-Tu. Translation assays were carried out with template encoding an N-terminal *myc*-tag followed by 15 consecutive amber stop codons, in the presence of L-tyrosine, D-tyrosine or the mixture of both, and the products were pulled down using anti-*myc* coated magnetic beads and analyzed by MALDI-TOF mass spectrometry (Figure 3a). With 8 μM EF-P, protein translations with *^L^*Tyr and *^D^*Tyr yielded up to 7 and 5 consecutive incorporations, while in the absence of EF-P, translations with *^L^*Tyr and *^D^*Tyr yield up to 3 and 2 consecutive incorporation respectively, based on the assumption that these peptides had similar ionization properties (Figure 3b). These results suggest that EF-P might have a role in facilitating translation in general, not limited to poly-proline sequences, as our result shows both poly-L and D-tyrosine peptide synthesis processivity are improved with the presence of EF-P.

**Figure 3.**
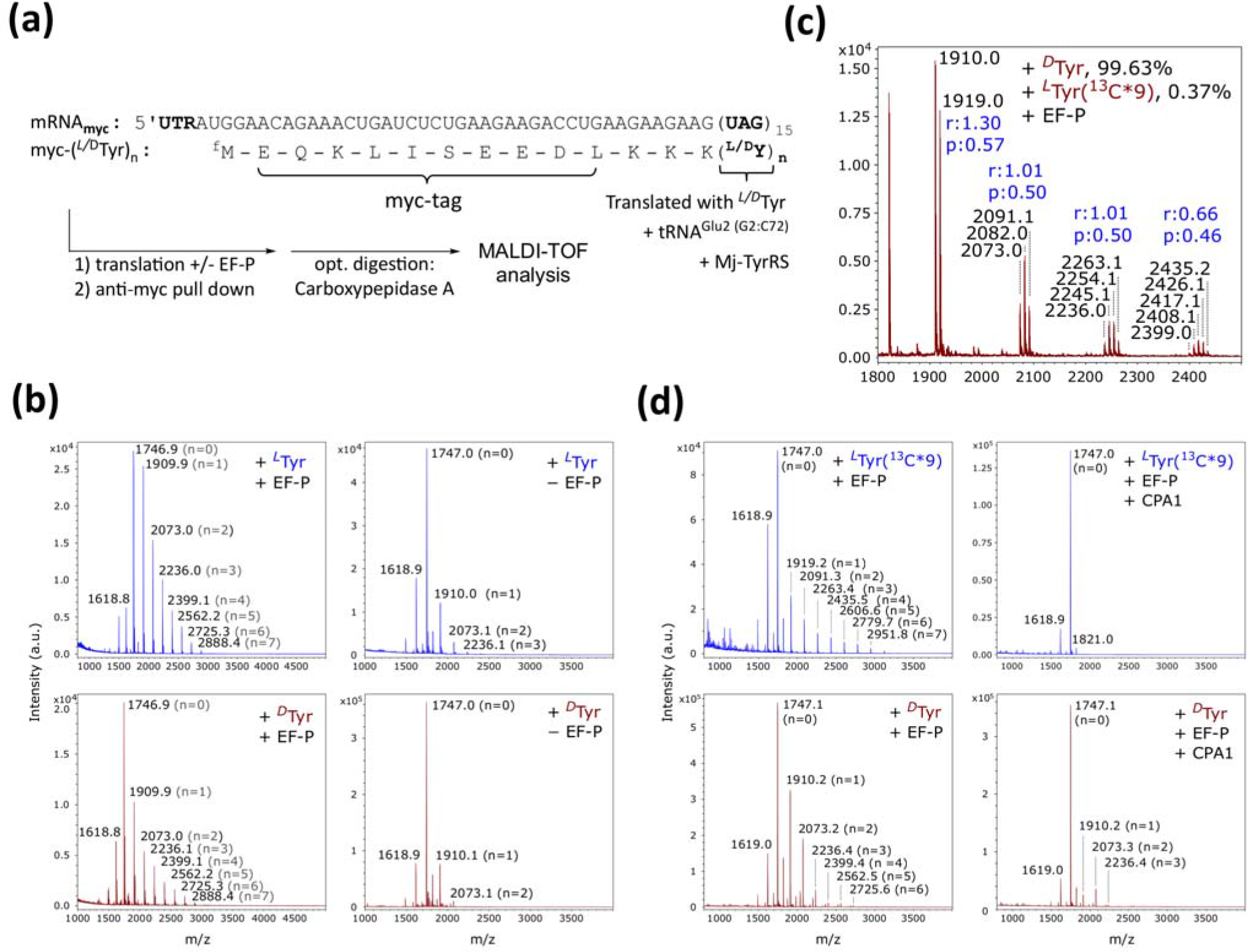
(a) Myc-peptide translation assay set up. The mRNA coding for N-terminus myc epitope is followed by three lysines and a stretch of 15 UAG codons. The resulting peptide products are optionally digested by bovine Carboxypeptidase A and analyzed by MALDI-TOF mass spectrometry. (b) MALDI-TOF-MS spectra of peptides from translation reactions with D- or L-Tyr, in the presence and absence of EF-P. Calculated mass for peptides myc-(*^L/D^*Y)_n_ are: 1745.92 (n=0), 1908.99 (n=1), 2072.05 (n=2), 2235.11 (n=3), 2398.18 (n=4), 2561.24 (n=5), 2724.30 (n=6), 2887.37 (n=7), whereas mass 1617.83 corresponding to ^f^Met-myc peptides followed by two lysines. (c) Zoom in MS spectra of peptides translated from *^D^*Tyr with 0.37% spiked in of *^L^*Tyr-^13^C*9. Incorporating every single *^L^*Tyr-^13^C*9 instead of *^L/D^*Tyr results in mass increase of 9.03 Daltons. “r” represents the intensity ratio of peptides with all *^D^*Tyr versus peptides with all *^L^*Tyr incorporation. “p” is the best fitted binomial distribution success probability of *^D^*Tyr incorporation, which are solved by the “Solve” function in Excel. (d) MS spectra of *^D^*Tyr and *^L^*Tyr peptides before and after carboxypeptidase A treatment. *^L^*Tyr peptides are all chopped up to n = 0 whereas C-terminal lysine halts further digestion, *^D^*Tyr peptides shows substantial resistance under the same condition. The original data and chi-square data are provided in Supplementary Table 3.

In crystal structures^18,22^, EF-P sits at the ribosome E-site and has close contact with P-site peptidyl-tRNA, suggesting that the P-site peptidy-tRNA might have a role in recruiting it. Comparing the sequences of all *E. coli* tRNAs, we noticed that proline tRNA and formylmethionine tRNA both carry a unique uridine insertion at position 17, which happens to be in close proximity to EF-P (Glu66, Thr67) according to a crystal structure (Supplementary Figure 1a, 1b). To test if this feature helps translation, we compared tRNA^Glu2^ with or without 17b uridine insertion in our myc-tyrosine peptide translation assay. However, we found that tRNA^Glu2^ with the uridine insertion gives no better or slightly less yields in carrying either L- or D-tyrosine into peptides (Supplementary Figure 1c). During the preparation of this manuscript, Katoh T. et al.^23^ reported that the entire 9-nt D-loop and a stable D-arm conserved in tRNA^Pro^ isoacceptors and tRNA^fMet^ is critical for the recruitment of EF-P. Given that the D-arm sequence of tRNA^Glu2^ is also unique among the 86 *E. coli* tRNAs which might contributes to its exceptionally high affinity to EF-Tu, further optimization of carrier tRNA would have to balance these two interacting partners.

To assess the degree of bias towards *^L^*Tyr versus *^D^*Tyr incorporation in this EF-P-ribosome translation system, in a reaction with *^D^*Tyr, we spiked in trace amount of ^13^C_9_-labeled *^L^*Tyr to assess the incorporation of *^D^*Tyr under competition. With 0.37% of ^13^C_9_-*^L^*Tyr in *^D^*Tyr, we observe ca. 1:1 ratio of mixed peptides with D- and L-tyrosine incorporation, and binomial distribution with the same p = 0.5 of mixed *^L/D^*Tyr incorporating products is observed for two, three and four consecutive tyrosine incorporations (Figure 3c). Because these distributions are only skewed slightly against consecutive *^D^*Tyr incorporations (Supplementary Table 3), suggesting that in our EF-P-ribosome system, peptide bond formation catalysis is uninfluenced by the underling amino acid’s chirality. Finally, to clear concerns regarding whether incorporations attributed to *^D^*Tyr were indeed actually *^D^*Tyr and not trace amounts of *^L^*Tyr impurity, we examined a bio-orthogonal property of these peptide products. Carboxypeptidase A selectively digests peptides from the C-terminus except for ionic residues, and the presence of *^D^*AAs at the P2, P1, and P1’ positions is known to greatly reduce the hydrolysis rate^24^. We therefore compared our myc-D/L-tyrosine peptides by MALDI-TOF analysis with and without Carboxypeptidase A digestion. As shown in Figure 3d, the peptide mixture translated with ^13^C_9_-labeled *^L^*Tyr is trimmed completely up to the C-terminal lysine, with no *^L^*Tyr left (m/z of [M+H]^+^ = 1746.92), while the peptide mixture translated with *^D^*Tyr exhibited substantial resistance under the same digestion condition. Aligned with what Schechter I. *et al.*^24^ found, we also observed semi-quantitatively that peptides with only one *^D^*Tyr at position P1’ are much less resistant to Carboxypeptidase A compared to peptides with two or three consecutive *^D^*Tyr in the P2, P1 and P1’ positions. Based on MALDI-TOF spectrum peak intensity, we estimated that 12% of peptide myc-(*^D^*Y)_1_ and 35% of myc-(*^D^*Y)_2_ survived digestion. Based on these data, we believe *^D^*Tyr has been incorporated into peptide through active ribosomal translation process.

In conclusion, we investigated important biases against *^D^*AA incorporation in protein synthesis. Our data demonstrated that the affinity between EF-Tu and amino acyl-tRNA plays a critical role in controlling *^D^*AA incorporation, and also that ribosome stalling on *^D^*AAs can be rescued by EF-P. Because EF-P also rescues stalling at poly-prolines, both forms of stalling may involve a common mechanism, and we hypothesize that the H-bond network required for the substrate-assisted mechanism of peptide bond formation is disrupted when an incorporated *^D^*AA or proline at the P-site is juxtaposed against a second instance of these residues at the A-site^25^. By combining the use of high affinity tRNA^Glu2^ and EF-P, we showed that the ribosome could polymerize chains of *^D^*Tyr as long as those of *^L^*Tyr. Our findings demonstrate that significant progress can be made in overcoming the bias against incorporating *^D^*AAs in translation at the ribosome, and suggest that the next key problem to address towards achieving D-protein synthesis will be to develop aaRSes that efficiently charge tRNAs with *^D^*AAs.

## Methods

Methods and any associated references are available in the online version of the paper.

## Acknowledgments

This work is sponsored by DOE grant DE-FG02-02ER63445, the Harvard Origin of Life Initiative, and EMD Millipore Inc. We thank Dr. Park Myung Hee at NIH for kindly sharing the plasmid pST39/His-EFP/YjeA/YjeK, and Mr. Jinfan Wang from Dr. Anthony Forster’s lab for discussion on tRNA 17A uridine insertion. We especially thank Dr. Jack Szostak, Dr. David Liu Dr. Jun Li and Dr. Dimitar Sasselov for valuable suggestions.

### Author contributions

P.H. conducted experimental design and execution with the help from F.W. and N.K., K.C. performed MALDI-TOF mass spectrum analysis, and J.A., R.S., S.T. and G.C. supervised the research. All of the authors contributed to writing the manuscript.

### Competing financial interests

The authors declare no competing financial interests.

### Additional information

Supplementary information and method is available in the online version of the paper. Reprints and permissions information is available online at http://www.nature.com/reprints/index.html. Correspondence and requests for materials should be addressed to G.C.

